# Automated extraction of electrode coordinates from structural MRI to assess tDCS placement accuracy

**DOI:** 10.64898/2025.12.08.693046

**Authors:** Sina Straub, Raphaela Schöpfer, Sarah Godehardt, Florian Wüthrich, Jessica Peter

**Affiliations:** University Hospital of Old Age Psychiatry and Psychotherapy, University of Bern, Bern, Switzerland; Translational Imaging Centre, Swiss Institute for Translational and Entrepreneurial Medicine, Bern, Switzerland; Graduate School for Health Sciences, Faculty of Medicine, University of Bern, Bern, Switzerland

**Author notes:** **Corresponding author:** Dr. rer. nat. Sina Straub University of Bern, Department of Old Age Psychiatry and Psychotherapy Bolligenstrasse 111, CH-3000 Bern 60.

**Keywords:** tDCS-MRI, electrode placement verification, neuromodulation

## Abstract

Accurate electrode placement is essential for studies using transcranial direct current stimulation (tDCS) since it determines whether - and to what extent - the intended brain region was modulated. Using in-scanner tDCS, placement accuracy can be determined by extracting electrode coordinates from structural MRI and by comparing actual and intended positions. Previous studies extracted coordinates manually, which is time-consuming and prone to error. Here, we present an algorithm to automate this process. The proposed algorithm first segments electrode and electrode-gel blobs visible on MRI. It then separates this segmentation into individual electrode regions and extracts their coordinates as the centroids of these regions. Finally, coordinates are assigned to the intended target positions using linear assignment. We evaluated the algorithm with T_1_-weighted MRI data from 65 individuals acquired at different scanners with different magnetic field strengths, image resolutions, and two high-definition montages by comparing the extracted coordinates with manual annotations from two independent readers. The median localization error of the algorithm was 2.4 mm (IQR 1.5 mm). The median variability between human readers was 2.7 mm (IQR 1.7 mm). Distances between algorithm-derived coordinates and the reference were smaller than distances between readers. Our proposed algorithm obviates the need for manual electrode segmentation and allows an objective evaluation of electrode placement accuracy for in-scanner tDCS. It may be used to relate electrode position errors to stimulation effects.

## 1 Introduction

Transcranial direct current stimulation (tDCS) is a non-invasive brain stimulation method that modulates the excitability of targeted brain regions by applying a weak constant current (typically 1 - 2 mA) to the scalp (Nitsche & Paulus, 2000; Stagg & Nitsche, 2011; Woods et al., 2016). It has been used to enhance or inhibit a wide range of cognitive functions, such as working memory (Fregni et al., 2005; Ruf et al., 2017), attention (Nikolin et al., 2019), or motor learning (Nitsche & Paulus, 2000). In addition, it is being investigated as an adjunct treatment for major depressive disorder (Brunoni, Ferrucci, et al., 2012; Yachou et al., 2025) or for promoting recovery after stroke (Brunoni, Nitsche, et al., 2012; Lefaucheur et al., 2017; Woods et al., 2016). tDCS can be applied using conventional large-pad montages, usually with two electrodes, or multi-electrode high-definition configurations (Müller et al., 2022) with small electrodes (most commonly in a 4 × 1 ring-shape). The latter enhance current focality and spatial accuracy in the target region (Nikolin et al., 2019). For both setups, electrode placement is commonly guided by cranial landmarks (Herwig et al., 2003; Jog et al., 2021; Okamoto et al., 2004; Rich & Gillick, 2019), although some studies used neuronavigation-based approaches (De Witte et al., 2018; Jog et al., 2021), which require multiple MRI scans. Deviations in electrode positioning alter current distribution patterns (Indahlastari et al., 2023; Niemann et al., 2024; Opitz et al., 2018; Woods et al., 2015) and the extent of cortical modulation. This likely contributes to variability in stimulation outcomes (Horvath et al., 2015; Indahlastari et al., 2019) and to reduced reproducibility across studies, particularly so for high-definition tDCS due to its higher focality. Hence, there is need for accurate and precise electrode placement.

When using in-scanner tDCS, structural MRI data can be used to extract the actual electrode positions. The distance from the intended position to the actual position is defined as the achieved placement accuracy. Quantifying electrode placement accuracy may help identify sources of error and reduce variability across studies. In addition, the actual electrode position can be related to the physiological, neuroanatomical, and behavioural effects of stimulation. In previous studies, electrode positions were extracted manually from structural MRI data (Indahlastari et al., 2021, 2023; Li et al., 2025; Niemann et al., 2024), a process that is time-consuming and may be prone to error or bias when performed by a single human reader. We therefore developed an automated algorithm to extract electrode positions (i.e., coordinates) from structural MRI data. We evaluated this algorithm using MRI data acquired with different MR scanners (3 T and 7 T), different head coils, different image resolutions and different high-definition electrode montages (two electrodes bifrontal or 4 × 1 ring-shaped). All montages targeted the left dorsolateral prefrontal cortex (dlPFC). We then validated the algorithm’s performance by comparing extracted electrode positions with manual annotations by two independent readers.

## 2 Material and methods

### 2.1 Participants

We included data of n = 65 participants of two randomised, double-blind, and sham-controlled clinical trials. Participants (mean age 31.1 ±16.3 years, age range 20-80 years, 38 female) were either healthy or had a diagnosis of major depressive disorder as confirmed by a clinical interview before study participation. All participants had to be fluent in German, right-handed, non-smokers, with normal or corrected-to-normal vision, no substance user disorder, and no history of neurological or psychiatric disorders (except for patients with depression). All data were collected at the University of Bern. All participants provided written informed consent before the study. The cantonal Ethics Committee approved the studies (2017-01664, 2024-01010) which were done in accordance with the Declaration of Helsinki and its amendments.

### 2.2 MRI data acquisition and processing

Participants received either sham or real high-definition tDCS (1 mA for 20 min; Soterix Medical Inc., Woodbridge, NJ, USA), while undergoing 3 T (MAGNETOM Prisma, Siemens Healthineers, Erlangen, Germany) (n = 15) or 7 T MRI (MAGNETOM Terra, Siemens Healthineers, Erlangen, Germany) (n = 50). We used a 1Tx-32Rx (Nova Medical Inc., Wilmington, MA, USA) head coil at 7 T or a 64 Rx head coil at 3 T. We acquired structural MRI data using a compressed-sensing accelerated MP2RAGE sequence with sagittal slice orientation (Table 1). During data processing, we generated structural head models using the automated pipeline CHARM, implemented in SimNIBS (Version 4.5) (Puonti et al., 2020; Thielscher et al., 2015). Each individual T_1_-weighted image served as input to the CHARM pipeline. Using the --forceqform flag ensured that the qform header information of the NIfTI file was used for spatial orientation. All other parameters were left at their default values.

**Table 1.** MP2RAGE sequence parameters.

|  | resolution | matrix size | repetition | echo | inversion | flip | bandwidth | acquisition |
| --- | --- | --- | --- | --- | --- | --- | --- | --- |
|  | in mm<sup>3</sup> |  | time | time | time | angle | in Hz/pixel | time |
|  |  |  | in ms | in ms | in ms | in ° |  | in min:sec |
| 7T | 0.63×0.63×0.63 | 384×384×256 | 6000 | 2.06 | 800/ 2700 | 4/ 5 | 240 | 7:40 |
| 3T | 1.00×1.00×1.00 | 256×256×176 | 5800 | 2.9 | 800/ 2200 | 5/5 | 240 | 4:14 |

*Abbreviations:* mm = millimetre, ms = milliseconds, Hz = Hertz, min = minutes, sec = seconds, T = Tesla.

### 2.3 Transcranial direct current stimulation

We used two different electrode montages and placed electrodes according to the 10-20 EEG system (Jurcak et al., 2007). For one set-up (4 × 1 ring-shaped), we placed the centre electrode (i.e., the anode) at position F3 and the four adjoining electrodes at F1, FC3, F5 and AF3 (n = 50). For the second set-up (bifrontal), we placed the anode at F3 and the cathode at F4 (n = 15). In total, this led to 280 individual electrodes to be extracted. We used silver-chloride ring electrodes or rectangular carbon rubber electrodes with corresponding electrode holders and EEG caps (all from Soterix Medical Inc.). We used paste (Abralyt, Easycap GmbH, Wörthersee-Etterschlag, Germany) and MR-visible gel (Signa gel, Parker Laboratories Inc., Fairfield, NJ, USA). We quantified the amount of MRI-visible gel in about two-thirds of participants.

### 2.4 Algorithm design

We developed an algorithm which was implemented in Matlab (Version R2024b, MathWorks, Inc., Natick, MA, USA). It uses the CHARM naming conventions for folder and file names (e.g., structural T_1_-weighted data and the tissue segmentation need to be stored as zipped NIfTI files “T1.nii.gz” and “final_tissues.nii.gz”). The proposed algorithm is freely available on github: <u>GitHub-SinaStraub/tDCS_automated_electrode_location_from_MRI</u>). The algorithm consists of electrode segmentation from structural MRI data (Step 1-4, Figure 1B-D) and coordinate extraction from this segmentation (Step 5-6; Figure 1E). Input parameters for our proposed algorithm are listed and described in Table 2.

**Figure 1:**
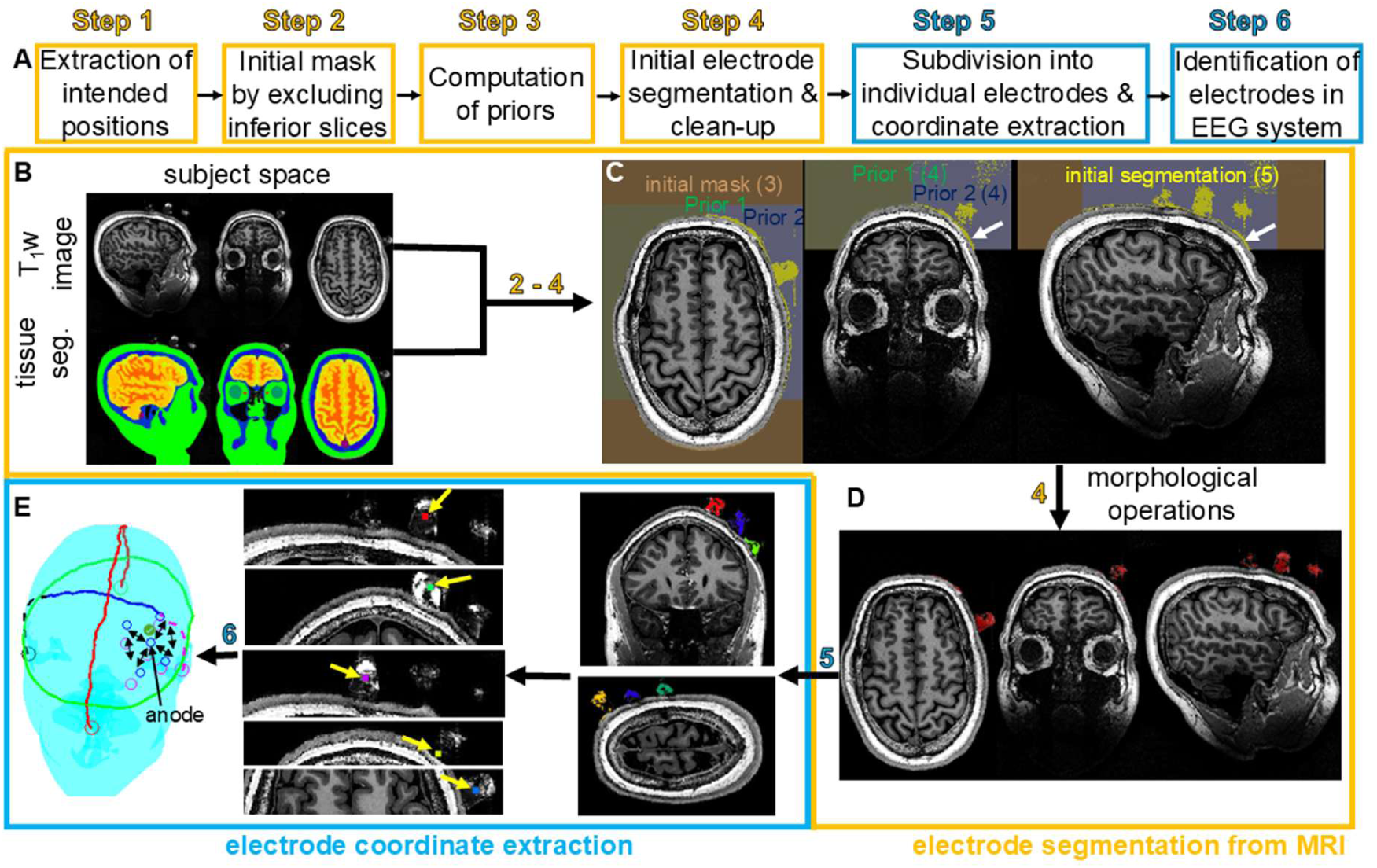
Flowchart of the algorithm (A) and graphical illustration of its’ six steps, including electrode segmentation (B-D) and coordinate extraction (E). (C) An initial mask (beige) includes all voxels above a certain slice threshold other than those segmented by CHARM’s tissue segmentation. Spatial priors (Prior 1 for the two electrode bifrontal montage, Prior 2 for 4 × 1 ring montage) were used to refine this initial mask. White arrows indicate voxels that were incorrectly segmented during this step. (D) Initial electrode mask (red). (E) Connected components into which the electrode mask was split and the extracted centroids (yellow arrows). At the end, a prompt opens that shows the actual (blue circles) and intended (magenta circles) electrode coordinates.

**Table 2.**
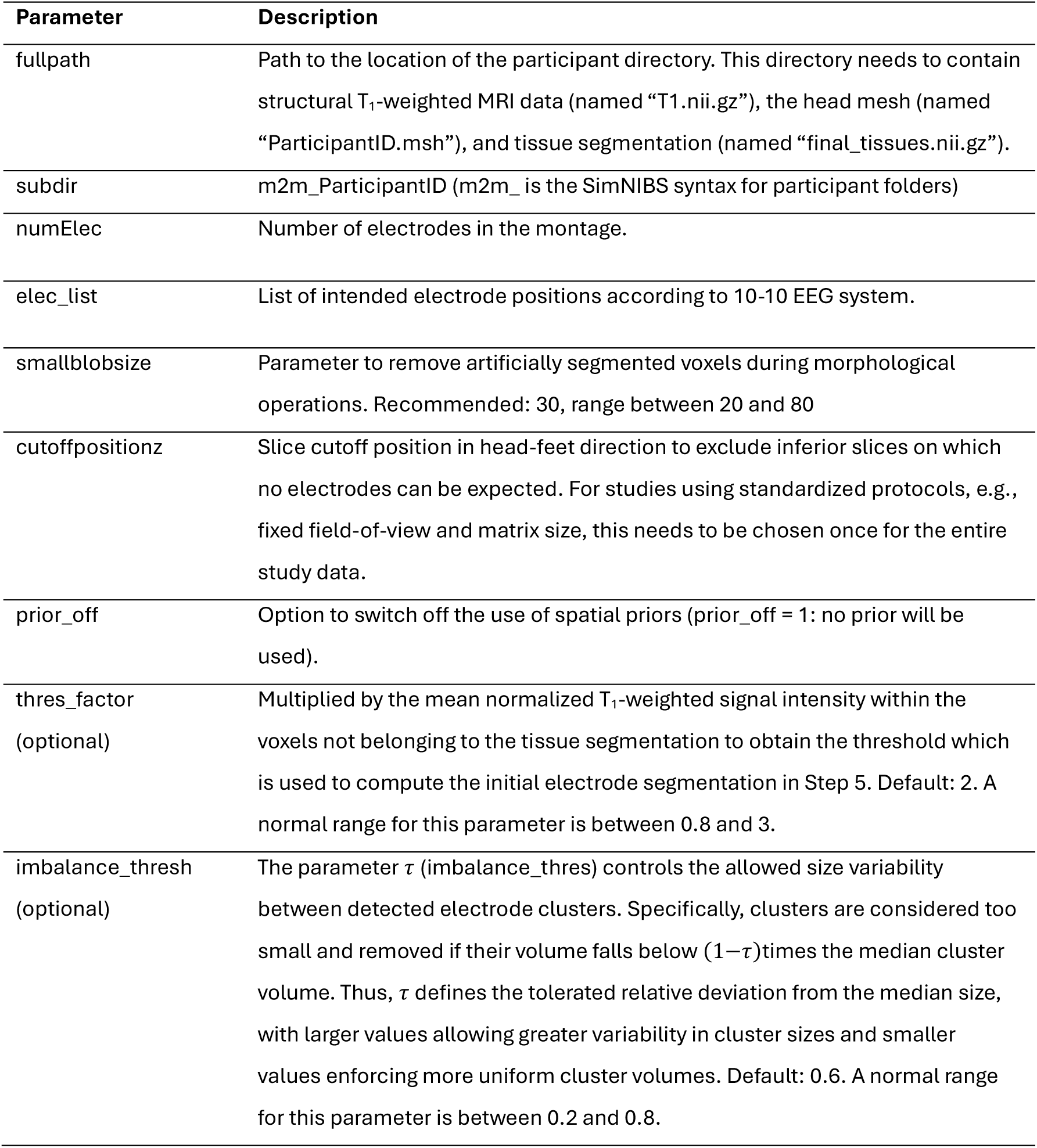
Input parameters for the proposed algorithm.

| Parameter | Description |
| --- | --- |
| fullpath | Path to the location of the participant directory. This directory needs to contain structural T<sub>1</sub>-weighted MRI data (named “T1.nii.gz”), the head mesh (named “ParticipantID.msh”), and tissue segmentation (named “final_tissues.nii.gz”). |
| subdir | m2m_ParticipantID (m2m_ is the SimNIBS syntax for participant folders) |
| numElec | Number of electrodes in the montage. |
| elec_list | List of intended electrode positions according to 10-10 EEG system. |
| smallblobsize | Parameter to remove artificially segmented voxels during morphological operations. Recommended: 30, range between 20 and 80 |
| cutoffpositionz | Slice cutoff position in head-feet direction to exclude inferior slices on which no electrodes can be expected. For studies using standardized protocols, e.g., fixed field-of-view and matrix size, this needs to be chosen once for the entire study data. |
| prior_off | Option to switch off the use of spatial priors (prior_off = 1: no prior will be used). |
| thres_factor<br>(optional) | Multiplied by the mean normalized T<sub>1</sub>-weighted signal intensity within the voxels not belonging to the tissue segmentation to obtain the threshold which is used to compute the initial electrode segmentation in Step 5. Default: 2. A normal range for this parameter is between 0.8 and 3. |
| imbalance_thresh<br>(optional) | The parameter $\tau$ (imbalance_thres) controls the allowed size variability between detected electrode clusters. Specifically, clusters are considered too small and removed if their volume falls below $(1-\tau)$ times the median cluster volume. Thus, $\tau$ defines the tolerated relative deviation from the median size, with larger values allowing greater variability in cluster sizes and smaller values enforcing more uniform cluster volumes. Default: 0.6. A normal range for this parameter is between 0.2 and 0.8. |

Step 1: The CHARM head mesh is used to extract coordinates of intended electrode positions based on the 10-10 EEG system (Jurcak et al., 2007). This is implemented in the function `get_standard_eeg_pos`. The voxel size of the data is extracted from NIfTI headers.

Step 2: An initial mask is computed by first excluding all voxels below a certain slice position where no electrodes can be expected. This slice position is an input parameter for the algorithm. Everywhere else, where the CHARM tissue segmentation is zero, this initial mask is set to one (Figure 1C).

Step 3: Based on the tissue segmentation, voxel size, and electrode configuration, a spatial prior is created to restrict electrode detection to anatomically plausible regions (Figure 1C): Specifically, boundary locations of the head are estimated along the superior–inferior (z), anterior–posterior (y), and left–right (x) directions by identifying the outermost non-zero voxels of the tissue segmentation. Using these boundaries, the search space is constrained by removing voxels outside predefined offsets from the scalp surface. These offsets are defined in physical units (millimetres) and converted to voxel space using the image resolution, ensuring consistency across data acquired with different imaging parameters such as image resolution or field of view. Depending on the electrode configuration (e.g., frontal left vs. bifrontal placements), additional lateral constraints are applied to exclude the contralateral hemisphere. This procedure results in a configuration-specific spatial mask that limits subsequent electrode detection to a plausible region of interest, thereby reducing false positives and improving robustness in challenging datasets. Although inter-individual variability in head size exists, the use of millimetre-based offsets provides a consistent approximation of electrode regions across subjects. Importantly, these constraints are not intended to localize electrodes directly but to restrict the search space. This aims to make the method robust to moderate anatomical differences. This step is implemented in the function `electrode_loc_prior` and can be omitted by setting the input parameter prior_of to one.

Step 4: After normalizing the T_1_-weighted image to a 0–1 intensity range, the mean voxel intensity within the resulting mask (Step 3) is computed. An initial electrode segmentation is obtained by retaining only voxels with normalized T_1_-weighted values exceeding this mean times a thresholding factor. This factor is provided to the algorithm as an input parameter (Table 2). The resulting initial electrode segmentation is then eroded and morphologically opened (Figure 1D).

Step 5: In this step, the function `correct_k_largest_clusters` cleans, splits, filters, and balances 3D clusters from the potentially merged or distorted initial electrode segmentation mask to reliably extract exactly K electrode centroids, where K is the number of electrodes in the montage. Connected components are first extracted from the binary electrode mask using 26-connectivity (Matlab function bwconncomp). For each component, volumetric size (number of voxels) and shape characteristics are computed using regionprops3. Elongation is quantified as elongation 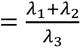, where 𝜆 ≥ 𝜆 ≥ 𝜆_ଷ_are the principal axis lengths. Components are classified as suspicious if they are either (i) highly elongated (elongation > 10) or (ii) unusually large, defined as having a volume greater than twice the mean volume of the K largest components. Suspicious components are subdivided using k-means clustering (k = 2), and resulting subcomponents are retained only if their elongation is below a stricter threshold (elongation < 8).

After this refinement, all valid components are reassembled and sorted by size. Small components are removed if their volume falls below (1−𝜏)times the median component size, where 𝜏 is a user-defined imbalance threshold. If fewer than K components remain, the largest remaining components are iteratively subdivided using k-means clustering (k = 2) until exactly K clusters are obtained. Finally, the centroid of each component is computed and used as the electrode position (Figure 1E).

Step 6: The anode and the cathodes are automatedly identified and assigned to corresponding EEG positions in a two-step procedure: First, the algorithm attempts to identify the anode by comparing voxel coordinates with the function `get_anode`. This is straightforward for the bifrontal montage but can become ambiguous for 4 × 1 montages. There, the algorithm sorts the x- and y-coordinates of all electrodes and identifies the anode as the electrode whose x and y values both fall in the middle of those sorted lists (Figure 1E). If this is successful, the anode is fixed. Second, detected electrode coordinates are automatedly assigned to a predefined EEG electrode layout (`assign_electrodes`) by minimizing spatial discrepancies between extracted and intended positions. EEG position coordinates are compared to detected coordinates using pairwise Euclidean distances in the horizontal (XY) plane. Optimal one-to-one assignments are then determined using `machtpairs`, the Matlab implementation of a linear assignment algorithm(Duff & Koster, 2001), ensuring a globally minimal assignment cost. Optionally, if the first electrode (e.g., the anode) has been fixed to a predefined position, it is excluded from reassignment. The resulting ordered electrode coordinates are saved in a text file in the participant folder and visualized for quality control. In theory, the second step alone would suffice, and it is also flexible enough to work for any electrode montage, however, for large deviations between actual and intended layouts, the two steps combined are more stable.

In addition, the algorithm creates a NIfTI file containing the centroids for each electrode (“landmarks.nii”) that allow the user to verify extracted positions on MRI data. Two NIfTI files containing the prior from Step 3 (“h1.nii” and “h2.nii”) are stored to help visualise which skull regions were used for electrode segmentation and to facilitate a quick adjustment of prior design (Step 3) for future electrode setups.

### 2.5 Algorithm implementation and evaluation

We used 270 as slice cutoff for 7 T data and 190 for 3 T data. We used 30 as the parameter ‘smallblobsize’, the default thresholding factor of two and 0.6 as imbalance threshold in the function that subdivides the binary electrode mask into individual electrodes (Table 2).

Algorithm evaluation and statistical analyses were performed in Matlab. To evaluate the accuracy of the algorithm, two independent readers (RS, FW; with three or eight years of neuroimaging experience) manually determined electrode locations from MRI data by placing electrode coordinates in the middle of the electrode ‘blobs’ using fsleyes (https://git.fmrib.ox.ac.uk/fsl/fsleyes/fsleyes/). These blobs corresponded to high-intensity regions on MR images from MR-visible gel surrounding the electrodes. We did not ask readers to differentiate between the anode and cathode(s) as this was deemed too difficult and time-consuming. Instead, we assigned the anode and cathodes automatedly in Matlab using the `assign_electrodes` function. If annotations between the two readers differed by more than 15 mm, readers were asked to reevaluate their annotations. The coordinates obtained by the two readers were averaged to define the reference coordinates, which were used to compute the distance between algorithm-derived coordinates and human annotations.

We quantified electrode localization accuracy between readers (i.e., inter-reader distance) as well as between readers and the algorithm (i.e., algorithm-reader distance) using Euclidean distances (median and interquartile range, IQR). The inter-reader distance also provided a reference for inter-rater variability. We visualised the distribution of distances using violin plots created with grpandplot (Wong, 2022). We then performed a pooled comparison between algorithm-reader and inter-reader distances using a one-sided Wilcoxon rank-sum test. Furthermore, Euclidean distances between the coordinates determined by the algorithm and the reader reference as well as inter-reader distances were analysed with a linear mixed-effects model fitted by restricted maximum likelihood (REML) using the `fitlme` function. The dependent variable was localization distance (in mm). Fixed effects included comparison type (algorithm–reader vs inter-reader), dataset (3 T ring, 7 T bifrontal, 7 T ring), and electrode polarity (anode vs cathode), as well as all interaction terms. To account for non-independence of electrodes within subjects, a random intercept for subject was included (y ∼ Method * Dataset * Polarity + (1|Subject)). Statistical significance was assessed using F-tests on marginal effects, with a significance level of p < 0.05 (two-tailed). To examine the relationship between inter-reader distance and algorithm-reader distance, we created a scatter plot.

To evaluate how deviations from intended electrode positions influenced the electric field, we performed electric field simulations using both intended and automatedly extracted electrode positions. We provide a detailed description in the Supplementary Material.

## 3 Results

Readers were asked to reassess their annotations for two participants as their annotations differed by more than 15 mm. For two participants, the default parameters of the algorithm had to be adjusted, as during visual quality control failed extraction of at least one electrode coordinate was identified (e.g. electrodes were not localized in ring shape, or one electrode was much further away from the others). For one participant, electrode visibility differed between electrodes (Figure 2A; adjusted parameters were thres_factor = 1.2 and imbalance_thres = 0.2). For the other individual, this was due to motion artifacts (Figure 2B; adjusted parameters were smallblobsize = 50 and thres_factor = 3). We used an average of 8.3 g of gel (7 T MRI data) or 5.7 g of gel (for 3 T MRI data). The readers reported difficulties in finding electrodes for the 3 T data, probably due to its lower spatial resolution and because less MR-visible gel was used.

**Figure 2:**
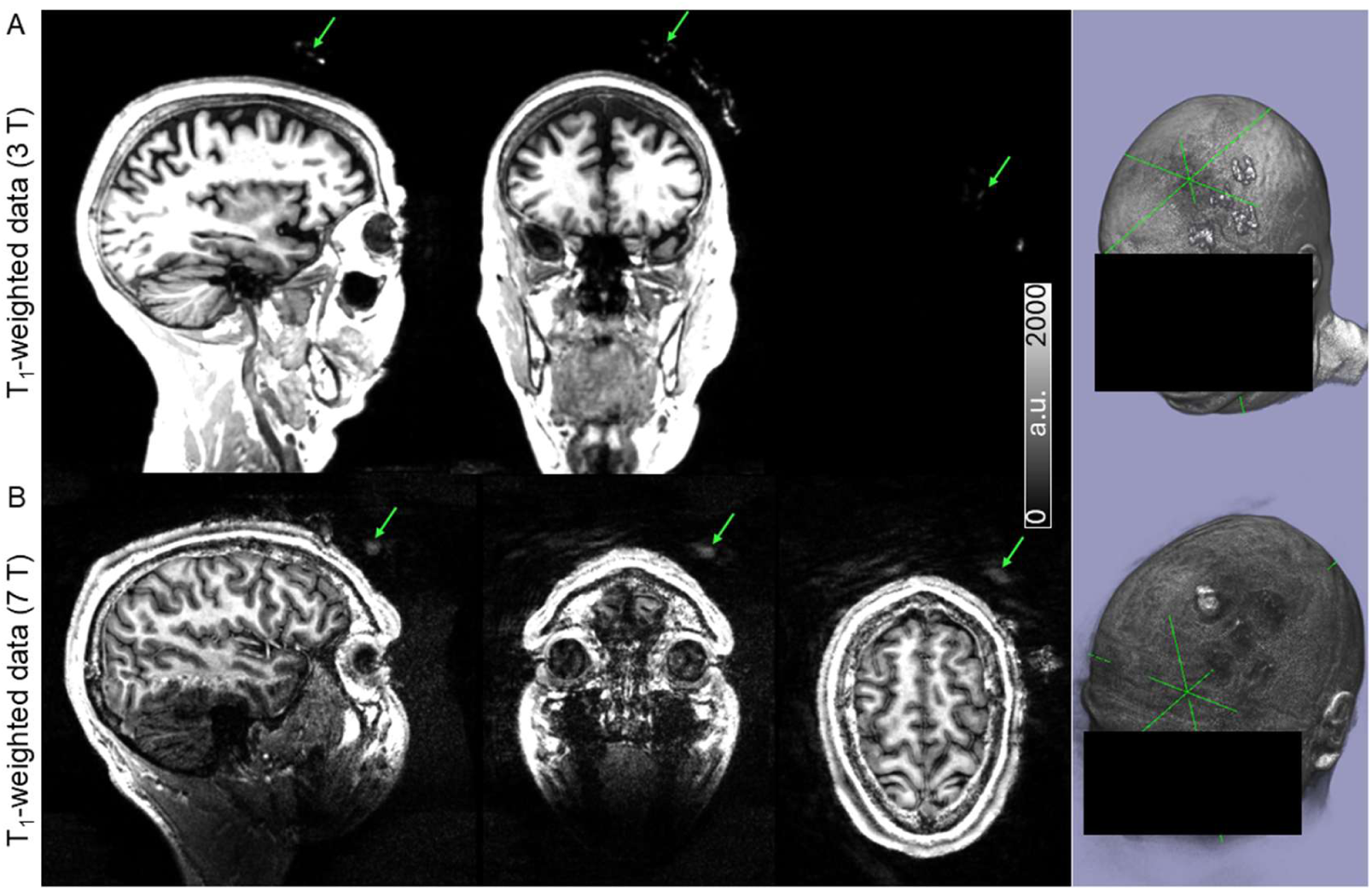
Example T_1_-weighted data acquired at 3 T (A) or 7 T (B). Example data show two participants for which default algorithm parameters needed to be adjusted due to disconnected electrode gel voxels (A, green arrow, green crosshair) or motion artifacts (green arrow, green crosshair)

Across all 280 electrodes, distances between readers ranged from 0.4 to 10.1 mm, while distances between algorithm and readers ranged from 0.4 to 9.9 mm. The median distance between algorithm-derived and reader-derived coordinates was 2.4 mm (IQR = 1.5 mm), which was slightly, but significantly, smaller than the median inter-reader distance of 2.7 mm (IQR = 1.7 mm, *p* < 0.001) (Figure 3). Across all datasets, the algorithm disagreed with human readers slightly less than the readers disagreed with each other (differences typically < 1 mm).

**Figure 3:**
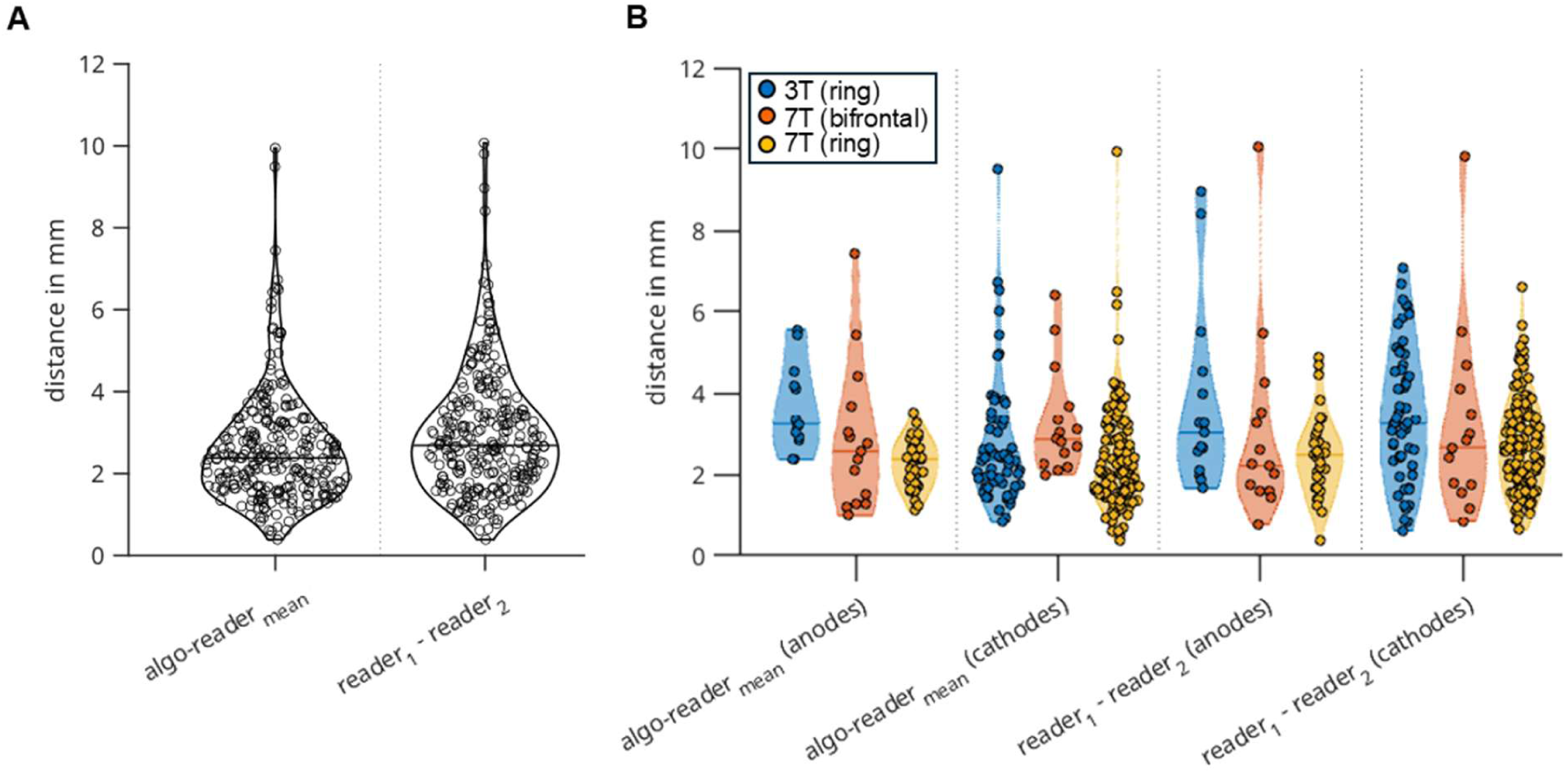
Distance between algorithm-derived coordinates and coordinates annotated by two readers. In addition, the distance between the two readers is shown. In Panel A data for all electrodes are pooled. In Panel B, data are grouped by scanner, electrode configuration (3 T – blue; 7 T two electrodes – orange; 7 T five electrodes – yellow).

The linear mixed-effects model revealed a significant effect of comparison type (F(1,548) = 7.57, *p* = 0.006), with algorithm-reader distances being smaller than inter-reader distances. There were also significant effects of dataset (i.e. scanner type and electrode placement - 3 T ring, 7 T bifrontal, 7 T ring) (F(2,548) = 3.19, *p* = 0.042) and electrode polarity (F(1,548) = 6.40, *p* = 0.012), indicating differences in localization distances across acquisition setups and between anodes and cathodes. A significant interaction between dataset and polarity was observed (F(2,548) = 3.51, *p* = 0.031), suggesting that polarity-related differences varied depending on the dataset. No significant interactions involving comparison type were found (all *p* ≥ 0.27), indicating that the difference between algorithm-reader and inter-reader distances was consistent across datasets and electrode polarities. The random intercept for subject (SD = 0.6 mm) indicates variability in average localization distances between subjects.

Only two electrodes showed algorithm-reader distances exceeding 8 mm, while there were four electrodes with inter-reader distances above 8 mm (Figure 4). Especially for the 3 T dataset, points tended to lie above the identity line, suggesting larger inter-reader distances relative to algorithm–reader distances (blue stars and dots).

**Figure 4:**
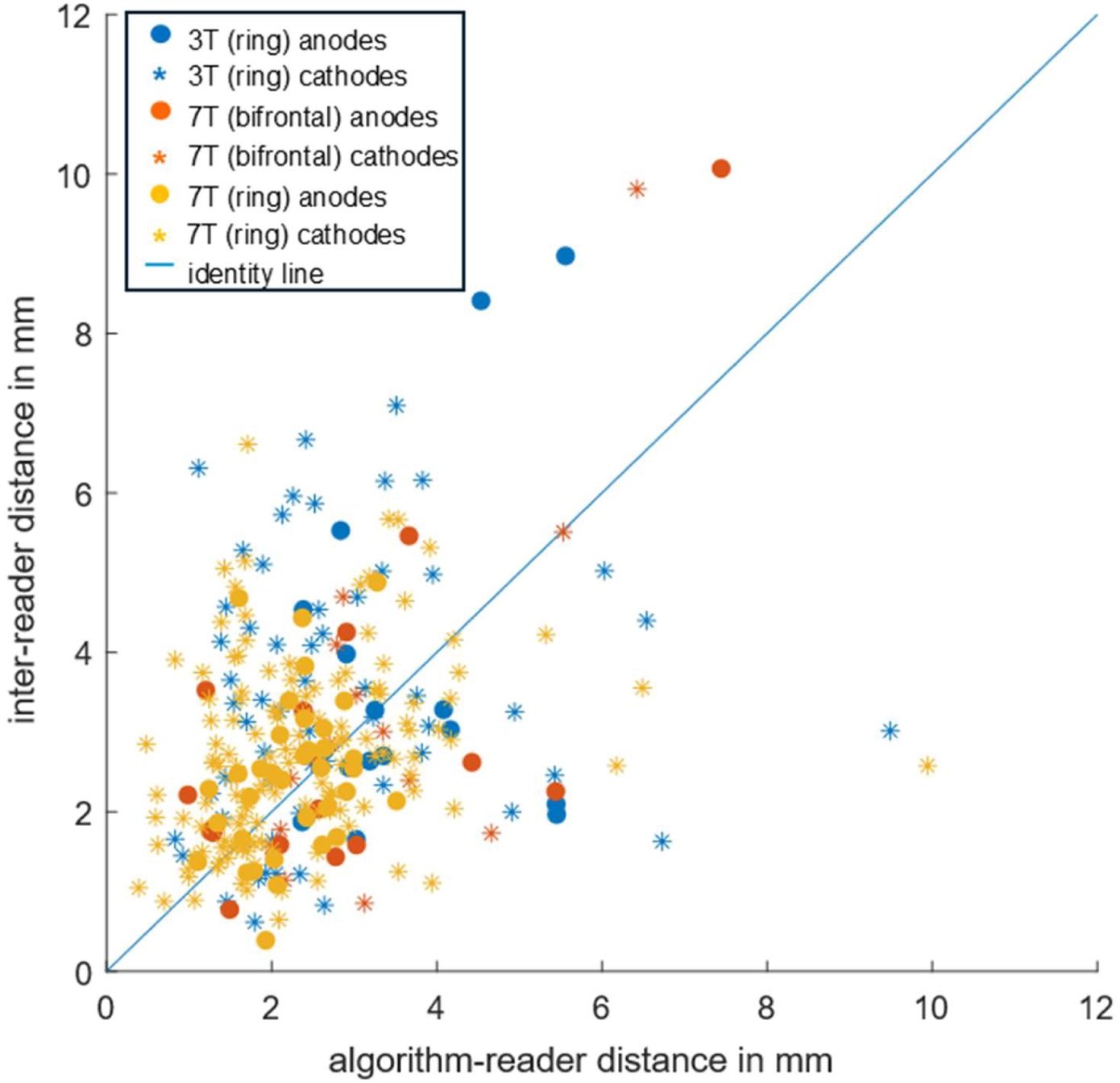
Scatter plot of inter-reader distance and algorithm-reader distance. The blue line indicates the identity line. Data are grouped by scanner and electrode configuration (3 T – blue; 7 T two electrodes – orange; 7 T five electrodes – yellow) and by cathodes (stars) and anodes (dots).

## 4 Discussion

This study introduces an algorithm that automatedly extracts tDCS electrode coordinates from structural MRI data. Both the pooled analysis and the linear mixed-effects model consistently showed a small (< 1 mm) but statistically significant difference between algorithm-reader and inter-reader distances, with slightly lower localization error for the algorithm, indicating that the algorithm performs at least at the level of human readers, with a slight advantage . The absence of interactions involving comparison type indicates that this difference was stable across datasets and electrode polarities. Although the magnitude of the difference varied numerically between conditions, as reflected in the distribution of data points (e.g., in the 3 T dataset), these variations did not translate into statistically significant interactions. The mixed-effects model further accounted for the non-independence of electrodes within subjects, providing a robust estimate of the overall difference between algorithm-reader and inter-reader distances across the full dataset. With a median localization error of 2.4 mm, the algorithm enables reliable and efficient quantification of electrode placement deviations. Given that an electrode placement accuracy of < 10 mm has been recommended for reliable stimulation (Opitz et al., 2018), the proposed algorithm meets the accuracy requirements for tDCS studies.

Our findings suggest that our proposed algorithm may obviate the need for manual extraction of electrode coordinates (Indahlastari et al., 2021, 2023; Niemann et al., 2024). While it cannot improve electrode placement itself, it provides a robust tool for quantifying the discrepancy between intended and actual electrode positions.

Adjustable parameters allow the algorithm to accommodate variations in signal intensity, spatial continuity, and electrode visibility (see Supplementary for a detailed discussion of algorithm design and specific parameters), ensuring consistent performance across datasets.

We deliberately evaluated the algorithm with data from two studies to demonstrate the flexibility and robustness of the proposed algorithm. It performed robustly across datasets acquired with different field strengths, head coils, image resolutions, and high-definition tDCS montages. We used commonly acquired T_1_-weighted MP2RAGE data, but the algorithm may be applied to other structural MRI data as long as tissue segmentation and head meshes (e.g., by using SimNIBS’ CHARM algorithm) are available, and electrode gel is visible on MRI. Because the algorithm excludes all tissue voxels using CHARM’s tissue segmentation, it does not rely on tissue-specific voxel intensities and is therefore largely insensitive to contrast variations introduced by acquisition parameters, field strength, or participant-specific factors such as age.

We used frontal montages in this study, but the algorithm can also be applied to posterior montages with minor adjustments. The 4 × 1 ring-shaped montage used here is particularly challenging for automated detection, as electrodes are placed very close together. Therefore, the algorithm is expected to perform even more accurately in configurations with fewer electrodes or wider spacing, such as 3 × 1 montages.

While electrode segmentations were used solely to extract coordinates in this study, segmentations of larger pad electrodes or MR-visible fiducial markers could also be used to estimate electrode orientations in future work. Deviations from intended electrode positions alter the electric field but do not directly predict stimulation efficacy. Such predictions require markers of target engagement, including cerebral blood flow (Baeken et al., 2017; Jog et al., 2021; Zheng et al., 2011), functional MRI (Antonenko et al., 2019; Jog et al., 2020; Li et al., 2025), or MR spectroscopy (Kim et al., 2014; Nandi et al., 2022). The algorithm proposed in our study can quantify the deviation between intended and actual electrode positions, which can then be linked to those markers. This provides a foundation for future studies to relate placement accuracy directly to physiological or behavioural effects.

Our study may have some limitations. First, the spatial priors were established for the specific electrode configurations of the current study. However, they can easily be adjusted to other electrode positions or target regions (e.g., by using additional elseif statements in the function `electrode_loc_prior`) or omitted (prior_of = 1). The validity of priors for other electrode montages has to be tested before it can be applied to larger studies. Second, the algorithm relies on the MR-visible gel surrounding the electrodes. It cannot be used for setups that do not use MR-visible gel or fiducial markers, and it may be more difficult to find electrodes reliably if the amount of MR-visible gel is low. Third, the algorithm has a few input parameter that need to be set empirically, and may differ from the ones used in this study. We have provided ranges for these input parameters.

In conclusion, the developed algorithm automatedly extracts actual electrode positions in studies using in-scanner tDCS with high accuracy. This obviates the need to manually segment electrode positions from structural MRI. The algorithm can be used to quantify deviations between actual and intended electrode positions, which can be linked to established markers of target engagement and, ultimately, to stimulation efficacy.

## Data and Code Availability

The code of the proposed algorithm is freely available on github (https://github.com/SinaStraub/tDCS_automated_electrode_location_from_MRI/tree/main). We also provide two datasets (bifrontal and 1 × 4 ring-shaped setup) to test the algorithm (https://doi.org/10.5281/zenodo.17860288).

## Author Contributions

S.S.: Conceptualization, methodology, formal analysis, visualization, writing-original draft, writing-review and editing; R.S.: Data acquisition, formal analysis, writing-review and editing; S.G.: Data acquisition, writing-review and editing; F.W.: formal analysis, writing-review and editing; J.P.: Conceptualization, writing-original draft, writing-review and editing, funding acquisition, supervision acquisition.

## Supporting information

Spplementary Material

## Acknowledgements

We thank Lea Raas, Aaron Friedli, Salome Flammer, Tamara Bachmann, Theresa Halbritter, Fabian Schmid and Jil Beckmann for their help in recruitment and data acquisition. We thank all participants for their time and support of the study.

