## Supplementary material for "Automated extraction of electrode coordinates from structural MRI to assess tDCS placement accuracy": Spplementary Material

### *1. Electric field simulations*

We computed the Euclidean distance between intended electrode positions and the positions extracted by the proposed algorithm. The latter, we call actual electrode position from here on. We used both actual and intended electrode positions to simulate how the observed deviations would affect the resulting electric field. Electric field simulations were performed with SimNIBS^1^ using CHARM’s head meshes. As input for SimNIBS, we entered electrode specifications (i.e., ring electrodes with outer diameter of 12 mm, inner diameter of 6 mm and a thickness of 3 mm), the use of 1 mA for stimulation, and used otherwise default parameters. Specifically, default electrode orientations were used. In SimNIBS, electrode centres are projected onto the scalp and the electrode normal is aligned with the local scalp surface normal, such that the electrode plane is tangential to the scalp. Because circular ring electrodes were used, the in-plane orientation is rotationally symmetric and does not influence the electric field distribution. Our code, as provided on github (<https://github.com/SinaStraub/tDCS_automated_electrode_location_from_MRI/tree/main>) includes a function simNibs_template for electric field simulations which partly reproduces code examples provided within SimNIBS. To quantify key features of the electric field magnitude distributions, we extracted the peak electric field coordinates and the 99.9^th^ percentile of the field magnitude from the subject-space electric field maps. In additions, electric field maps were computed in fsaverage space^2,3^. Using the latter, we generated group-level maps of electric field magnitude mean and standard deviation. To assess the electric field in the target region (i.e., dlPFC), we used the labels p9-46v, 46, and 9-46d from the HCP_MMP1.0 parcellation in fsaverage space^2,3^. These regions are located within the dlPFC and roughly correspond to Brodmann areas 9 and 46. We also considered other labels in the vicinity of the dlPFC (a9-46v, IFSa, 8Ad, 8Av, 8BL, 8C, s6-8, i6-8, 9a, 9p) that correspond to Brodman areas 8, 9, 46 or to transitions zones between them (except for IFSa, the anterior part of the inferior frontal sulcus). Distributions of subject-level mean region-of-interest (ROI) electric field magnitude are shown using violin plots created with grpandplot^4^ and across subjects median E-field magnitudes with IQRs are reported. We assessed whether the simulated electric field magnitudes were significantly smaller for actual compared to intended electrode position using one-sided Wilcoxon signed-rank tests. *p* < 0.05 was considered statistically significant. For visualization, we converted electric field peak coordinates to MNI space using subject2mni_coords.

*2. Assessment of E-field variations induced by deviations from intended electrode positions*

Distances between intended and actual electrode positions ranged from 4.3 mm to 51.0 mm (median = 21.7 mm, IQR = 10.8 mm).

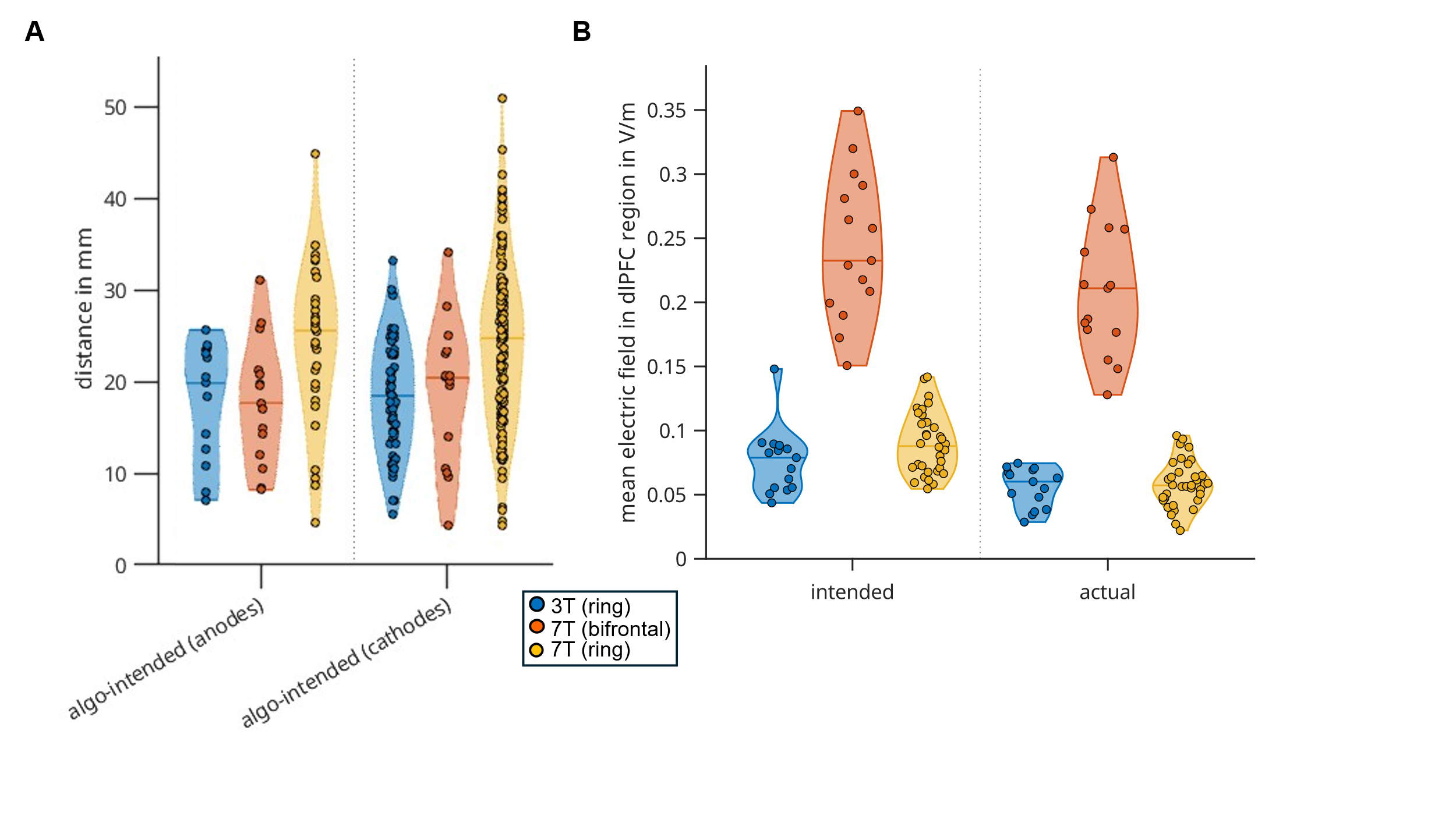

**Figure S1:** (A) Violin plots show the distances between the electrode coordinates extracted by the algorithm (actual) and the intended locations. (B) Mean electric field magnitude within the dlPFC for intended and actual electrode positions. Data are grouped by scanner and electrode configuration (3 T, blue; 7 T, bifrontal - orange; 7 T, ring-shaped - yellow).

We found significantly lower mean electric field magnitude for intended compared to actual electrode positions in the target region for all data groups and setups (Figure S1B, Table S2). IQRs were more than twice as large for the bifrontal setup (0.08 – 0.09 V/m) than for the ring-shaped setup (0.02 – 0.04 V/m). For the bifrontal setup the median anode distance from the intended positions was 17.7 mm and the median electric field reduction in the dlPFC ROI was about 7 %. It was about 33 % for the ring setup from the 7 T data for which median anode distance was 25.6 mm, and 25 % for the ring setup from the 3 T data, for which median anode distance was 19.9 mm.

**Table S2.** Mean electric field strength in the dlPFC. p-Values represent results of Wilcoxon signed-rank tests (one-sided). Significant values are shown in bold.

|  | intended | |  | actual | |  |
| --- | --- | --- | --- | --- | --- | --- |
| field strength and set-up | median | IQR |  | median | IQR | *p*-value |
| 7 T, bifrontal | 0.23 | 0.09 |  | 0.21 | 0.08 | **< 0.001** |
| 7 T, ring-shaped | 0.09 | 0.04 |  | 0.06 | 0.02 | **< 0.001** |
| 3 T, ring-shaped | 0.08 | 0.03 |  | 0.06 | 0.03 | **< 0.001** |

*Abbreviation*: IQR = interquartile range.

These observations are likewise reflected in the group-averaged electric field magnitude maps (Figure S3), which exhibit a higher standard deviation when based on the actual electrode coordinates compared to the intended coordinates. This indicates that electrode placement deviations reduced the overall group-averaged electric field strength and increased variability between participants.

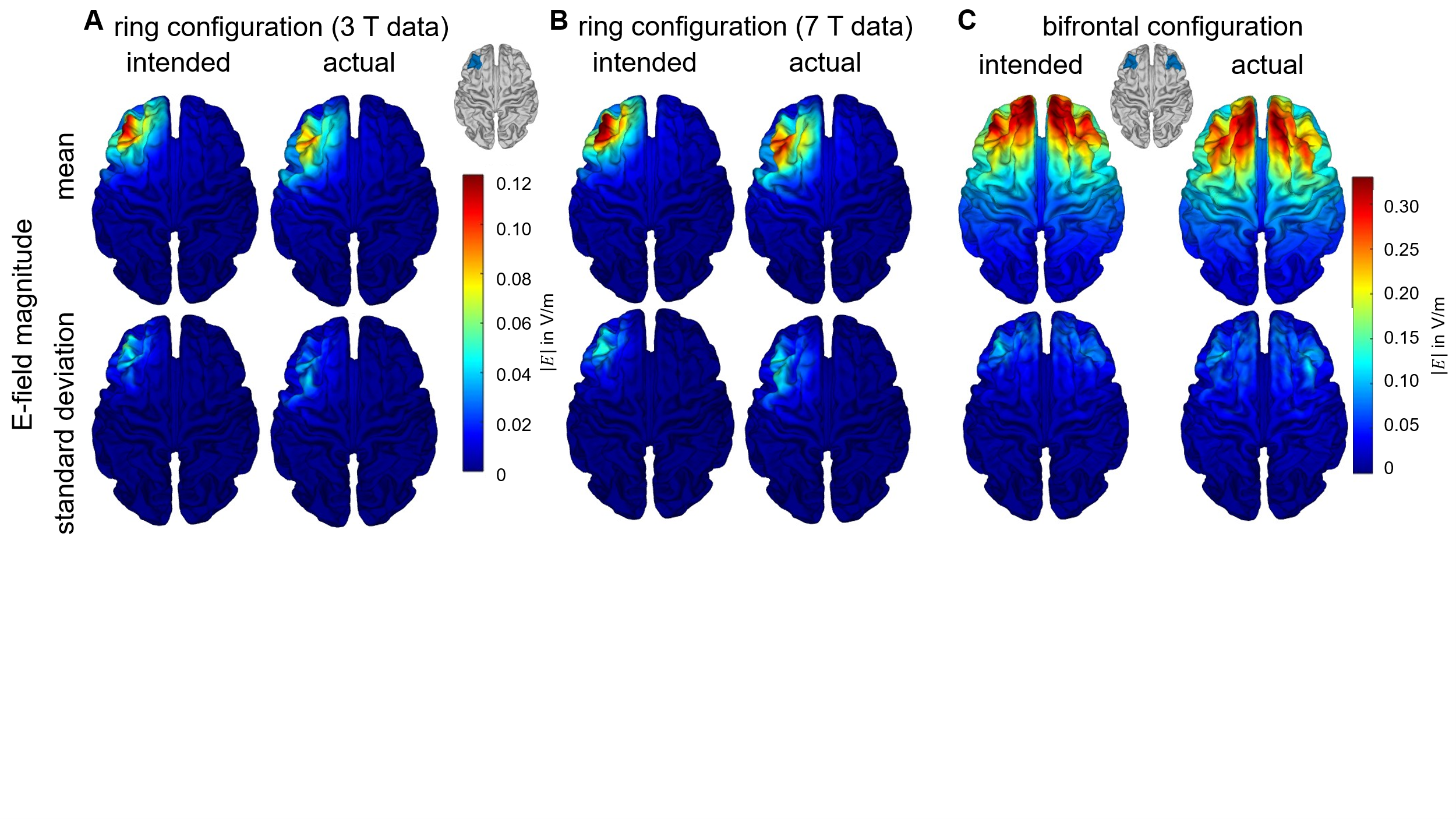
**Figure S3:** Maps of mean and standard deviation of electric field magnitude for all participants (simulated for intended and actual electrode positions).

Because the electrodes were not placed exactly as intended (Figure S4, see also Figure S1A), the coordinate of the peak electric field shifted noticeably. With the intended placements, around 50 % of bifrontal and 95 % of 4 × 1 ring montage peak field coordinates laid within the dlPFC. However, when using the actual electrode positions (Figure S4A), the peak field coordinates more frequently occurred outside the dlPFC (Figure S4B) - over 50 % of cases in the bifrontal setup and over 60 % in the ring setup.

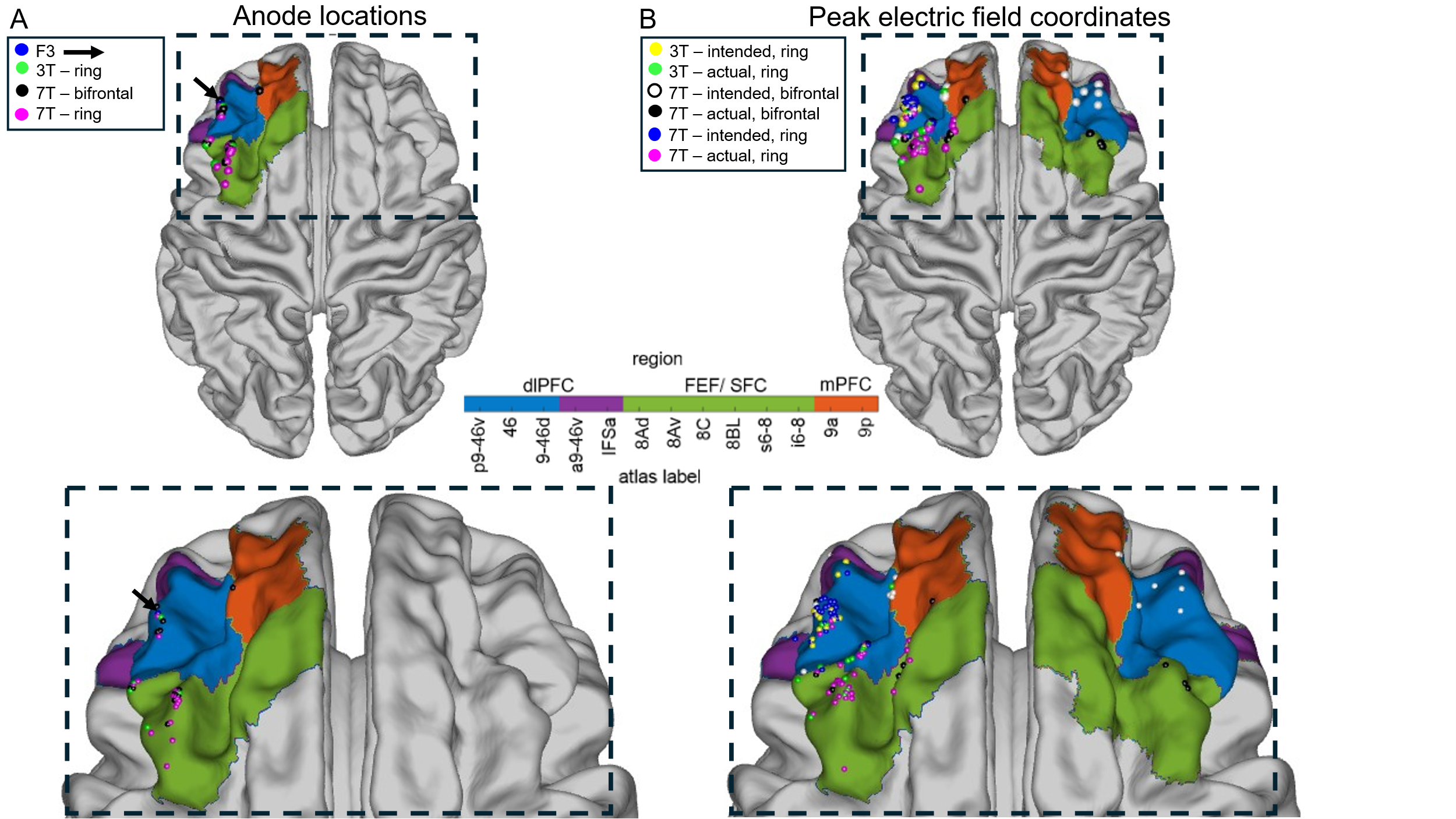

**Figure S4:** Intended (blue) and actual (pink, green, yellow) anode locations (A) as well as resulting peak electric field coordinates (intended yellow, white, blue; actual green, black, pink) (B) overlaid on fsaverage cortical surface. Colours indicate the dorsolateral prefrontal cortex (dlPFC), the frontal eye field / superior frontal cortex (FEF/ SFC), and the medial prefrontal cortex (mPFC). Labels were used from the HCP_MMP1.0 atlas.

*3. Algorithm design*

MR images may contain artifacts - for example, due to motion - that appear as noisy, high-intensity regions resembling the signal of gel-filled electrode holders. To mitigate this, we incorporated a cutoff in the head–feet direction and a spatial prior during segmentation to constrain plausible electrode locations (e.g., left/right hemisphere or anterior/posterior regions). Conversely, when little gel is used, electrodes may produce only a weak MR signal and appear faint or fragmented. To accommodate such variability in signal intensity and spatial continuity, the algorithm uses adjustable input parameters. In particular, a thresholding factor and the parameter ‘smallblobsize’ adapt the initial electrode segmentation to differences in visibility and spatial extent. The subsequent coordinate extraction step, in which the segmentation is subdivided into individual electrodes, is designed to remain robust even under poor visibility conditions.

The parameter $\tau$(imbalance threshold) can be selected based on the expected variability in electrode cluster sizes. Lower values (e.g., $\tau\approx0.1$–0.2) enforce stricter size uniformity and are appropriate to exclude smaller clusters. The latter is more important when little gel was used or when gel blobs appear somewhat disconnected. Higher values (e.g., $\tau\approx0.3$–0.8) allow for greater variability and are preferable when some electrodes are expected to appear smaller due to partial volume effects, lower signal intensity, or imperfect segmentation.

1.	Saturnino GB, Puonti O, Nielsen JD, Antonenko D, Madsen KH, Thielscher A. SimNIBS 2.1: A Comprehensive Pipeline for Individualized Electric Field Modelling for Transcranial Brain Stimulation. In: Makarov S, Horner M, Noetscher G, eds. *Brain and Human Body Modeling*. Cham: Springer International Publishing; 2019:3-25. doi:10.1007/978-3-030-21293-3_1

2.	Glasser MF, Coalson TS, Robinson EC, et al. A multi-modal parcellation of human cerebral cortex. *Nature*. 2016;536(7615):171-178. doi:10.1038/nature18933

3.	Petre B, Ceko M, Friedman NP, et al. 2016 Glasser MMP1.0 Cortical Atlases. 2023:534271010 Bytes. doi:10.6084/M9.FIGSHARE.24431146.V8

4.	Wong MH. grpandplot: An open-source MATLAB tool for drawing box plot and violin plot with automatic multi-way data grouping. November 2022. doi:10.5281/ZENODO.7295877
